# The dynamics of centromere assembly and disassembly during quiescence

**DOI:** 10.1101/2025.09.08.674938

**Authors:** Océane Marescal, Kuan-Chung Su, Brittania Moodie, Noah J. L. Taylor, Iain M. Cheeseman

## Abstract

Quiescence is a state in which cells undergo a prolonged proliferative arrest while maintaining their capacity to reenter the cell cycle. Here, we analyze entry and exit from quiescence, focusing on how cells regulate the centromere, a structure involved in chromosome segregation. Despite the constitutive localization of centromere proteins throughout the cell cycle, we find that cells rapidly disassemble most centromere proteins during quiescence entry, while preserving those required to maintain centromere identity. During quiescence exit, the centromere is reassembled and rapidly regains normal homeostatic levels of centromere proteins. Although the histone variant CENP-A is typically deposited during G1, we find that CENP-A deposition does not occur during the G1 immediately following quiescence exit, and instead occurs after cells complete their first mitosis. In contrast, other centromere proteins relocalize during the first S phase independent of DNA replication. These findings reveal centromere dynamics during quiescence entry and exit and highlight paradigms for the timing and control of centromere protein deposition.

## Introduction

Quiescence is a state of reversible proliferative arrest in which cells are no longer dividing but retain the capacity to reenter the cell cycle^1^. Quiescent cells play diverse functions and are widespread in an organism. Examples include hepatocytes, oocytes, and tissue-resident stem cells, all of which can remain in a non-dividing state for years, only reentering the cell cycle when activated with the appropriate stimulus^1–11^. Thus, quiescent cells are faced with the unique task of preventing division while simultaneously preserving their capacity to divide. This need to achieve a balance between growth arrest and the capacity for cell cycle reentry is reflected in the numerous changes to gene expression and metabolism that characterize the quiescent state^1,12–23^. For example, quiescent cells downregulate the expression of genes that promote cell cycle progression but upregulate those involved in protecting the cell from long-term DNA damage^12,13,17,18^. Quiescent cells also undergo large-scale alterations to cellular structures and organelles, including the rearrangement of centrioles to form primary cilia^24–27^, a decrease of nuclear pore size and distribution^28–30^, and a reduction of mitochondrial number^31–33^.

One key cellular structure that represents an important consideration for quiescent cells is the centromere. During mitosis, the centromere recruits the macromolecular kinetochore complex, which in turn is crucial for mediating microtubule attachments and segregating the genetic material^34^. Defining this region of the chromosome does not occur through specific DNA sequences in vertebrate cells. Instead, centromeres are specified epigenetically through the presence of the centromere-specific histone H3 variant, CENP-A^34–39^. Previous work from our lab found that quiescent cells slowly, but continuously incorporate new CENP-A nucleosomes at centromeric regions^40^. This continuous CENP-A deposition maintains centromere identity throughout an extended quiescent arrest and preserves the cell’s ability to segregate its chromosomes upon cell cycle reentry. In addition to CENP-A, 16 other CENP proteins localize constitutively to the centromere in all phases of the cell cycle in proliferating cells^41^. These proteins, collectively known as the constitutive centromere-associated network (CCAN), can be divided into 5 groups that are interdependent on each other for localization: CENP-C, the CENP-L/N complex, the CENP-H/I/K/M complex, the CENP-O/P/Q/U/R complex, and the CENP-T/W/S/X complex^34,41^. CCAN components serve to recruit key microtubule-binding proteins to the chromosomes during mitosis, thus forming the centromere-kinetochore interface^42–47^. The fate of these CCAN components as cells enter and exit quiescence remains poorly understood.

Here, we investigate the behavior and regulation of centromere components during quiescence entry and exit. Despite their constitutive localization in cycling cells, we find that centromere proteins are rapidly lost upon quiescence entry. This centromere disassembly is regulated at the level of transcription through the downregulation of most CCAN components. In contrast, we show that CENP-C is uniquely preserved at low levels in quiescent cells where it contributes to CENP-A deposition. Finally, we analyze the dynamics of centromere reassembly during cell cycle reentry and exit from quiescence. We find that most CCAN proteins are re-deposited at centromeres during the S phase immediately following quiescence exit in a process that occurs independently of DNA replication. This unique *de novo* assembly event provides a novel situation to evaluate centromere protein dynamics and behavior. Conversely, new CENP-A is not deposited until after the first mitotic division following quiescence release. This CENP-A deposition event re-equilibrates centromeric CENP-A back to the steady state levels observed in cycling cells. This study defines centromere protein behavior and dynamics during quiescence and highlights how quiescent cells can concurrently disassemble a structure that is not necessary during growth arrest, while preserving certain components to enable cell cycle reentry.

## Results

### The centromere is rapidly disassembled upon quiescence entry

To analyze the behavior of centromere proteins in quiescent cells, we induced non-transformed human retinal pigment epithelial (RPE1) cells to enter quiescence using a combination of serum starvation and contact inhibition. This treatment successfully induced quiescence, as monitored by lack of EdU incorporation and 2n DNA content (Figure S1A, B, C). Following quiescence induction, we used immunofluorescence to monitor the localization of CCAN proteins over the course of 7 days. The localization of the centromeric histone H3 variant, CENP-A, persisted during quiescence (Figure S1D), as described previously^40^. In contrast, the rest of the centromere CCAN proteins were rapidly disassembled upon entry into quiescence (Figure 1A-J). In particular, CENP-T, CENP-L, and CENP-K showed a marked reduction of centromere intensity within just 24 hours of quiescence induction and were completely lost after 5 days (Figure 1A-F). CENP-O/P levels remained stable for a longer period but were eventually lost by 5 days of quiescence (Figure 1G, H). Although CENP-C centromere intensity also decreased in quiescent cells, its levels plateaued to around one-fourth of that found in cycling cells, maintaining diminished, but stable centromere localization during quiescence (Figure I, J). This pattern of quiescence CCAN behavior was conserved in mouse 3T3 cells (Figure S1E, F, G, H), which also showed CENP-T loss following quiescence entry, but CENP-C retention.

**Figure 1:**
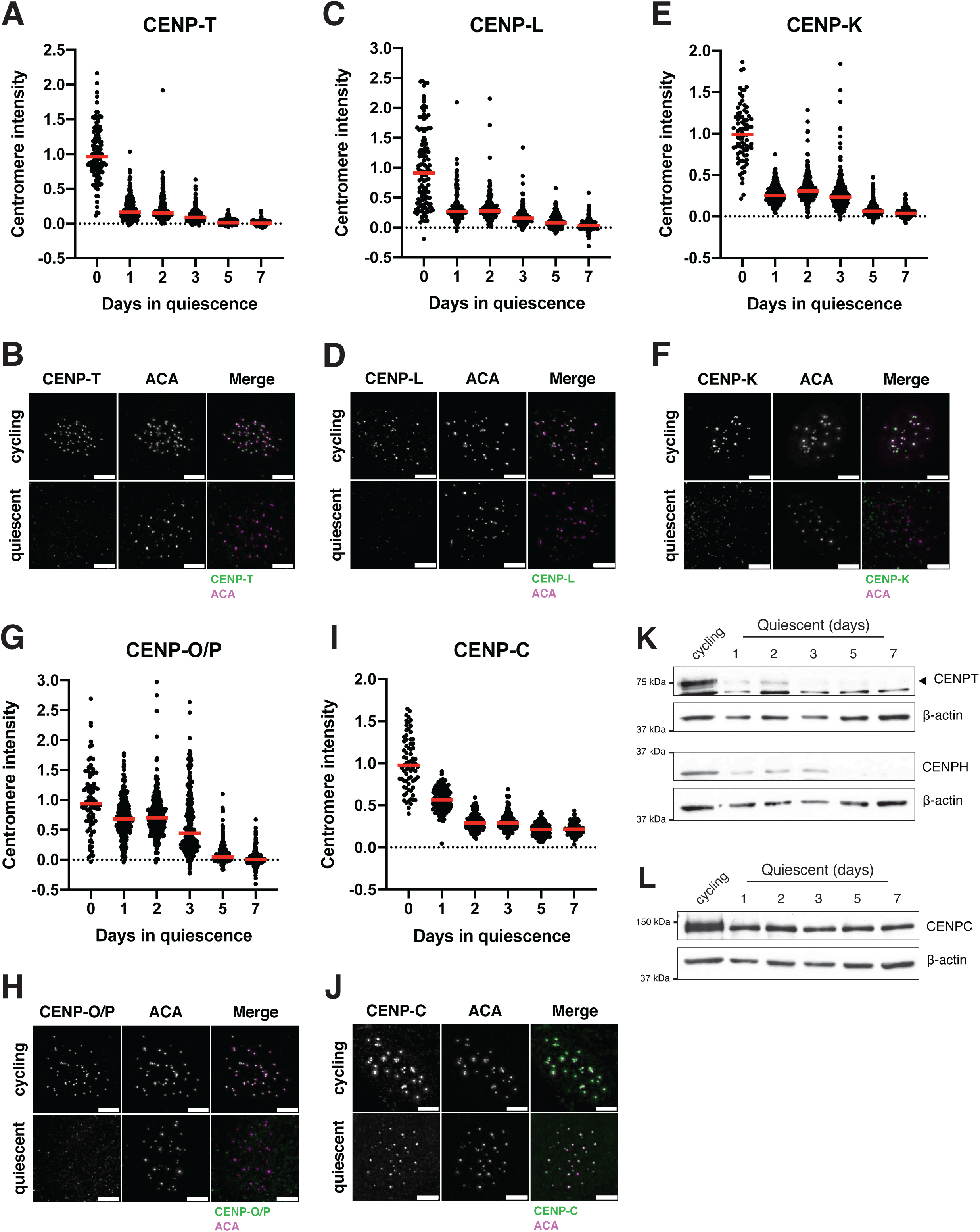
The centromere is rapidly disassembled upon quiescence entry. A. Graph showing CENP-T centromere intensity level over time of quiescence entry. Each point indicates the average centromere intensity level for all centromeres in a single cell, adjusted for background. Intensity values were normalized to day 0. Red line represents the median. Points were aggregated from 2 replicates. n = 122, 262, 255, 261, 258, and 271 cells for 0, 1, 2, 3, 5, and 7-day time points respectively. B. Representative immunofluorescence images of a cycling and quiescent cell. Cells were stained with CENP-T and anti-centromere (ACA) antibodies. Scale bar = 5µm. C. Graph showing CENP-L centromere intensity level over time of quiescence entry. Each point indicates the average centromere intensity level for all centromeres in a single cell, adjusted for background. Intensity values were normalized to day 0. Red line represents the median. Points were aggregated from 2 replicates. n = 120, 166, 178, 156, 156, and 160 cells for 0, 1, 2, 3, 5, and 7-day time points respectively. D. Representative immunofluorescence images of a cycling and quiescent cell. Cells were stained with CENP-L and anti-centromere (ACA) antibodies. Scale bar = 5µm. E. Graph showing CENP-K centromere intensity level over time of quiescence entry. Each point indicates the average centromere intensity level for all centromeres in a single cell, adjusted for background. Intensity values were normalized to day 0. Red line represents the median. Points were aggregated from 2 replicates. n = 80, 302, 343, 407, 324, and 242 cells for 0, 1, 2, 3, 5, and 7-day time points respectively. F. Representative immunofluorescence images of a cycling and quiescent cell. Cells were stained with CENP-K and anti-centromere (ACA) antibodies. Scale bar = 5µm. G. Graph showing CENP-O/P centromere intensity level over time of quiescence entry. Each point indicates the average centromere intensity level for all centromeres in a single cell, adjusted for background. Intensity values were normalized to day 0. Red line represents the median. Points were aggregated from 2 replicates. n = 88, 291, 285, 266, 254, and 252 cells for 0, 1, 2, 3, 5, and 7-day time points respectively. H. Representative immunofluorescence images of a cycling and quiescent cell. Cells were stained with CENP-O/P and anti-centromere (ACA) antibodies. Scale bar = 5µm. I. Graph showing CENP-C centromere intensity level over time of quiescence entry. Each point indicates the average centromere intensity level for all centromeres in a single cell, adjusted for background. Intensity values were normalized to day 0. Red line represents the median. Points were aggregated from 2 replicates. n = 74, 144, 140, 114, 140, and 93 cells for 0, 1, 2, 3, 5, and 7-day time points respectively. J. Representative immunofluorescence images of a cycling and quiescent cell. Cells were stained with CENP-C and anti-centromere (ACA) antibodies. Scale bar = 5µm. K. Western blots of cells in quiescence for the indicated amount of days. Blots were incubated with CENP-T and CENP-H antibodies. β-actin is used as a loading control. L. Western blot of cells in quiescence for the indicated amount of days. Blot was incubated with CENP-C antibody. β-actin is used as a loading control.

In addition to measuring centromere localization, we conducted western blotting to monitor centromere protein levels (Figure 1K, L, Figure S1I, J, K). CENP-T and CENP-H proteins were depleted in quiescent cells, whereas CENP-C protein was retained, consistent with their localization behavior. Thus, centromere components are rapidly disassembled upon quiescence entry through the depletion of most centromere components, whereas CENP-C is retained with reduced centromere levels.

### Quiescent cells regulate centromere behavior through a transcriptional program

To determine how centromere disassembly is regulated, we next conducted RNA sequencing in quiescent and cycling cells (Figure 2A, Figure S2A). We observed a strong decrease in the expression of most centromere protein mRNAs in quiescent cells, with the exception of CENPC mRNA, which was increased by around 2-fold. We additionally verified these results using qPCR (Figure 2A, B, C). We also observed similar trends in mouse 3T3 cells (Figure S2B). These changes in mRNA levels occurred within 24 hours of quiescence induction, closely mirroring the dynamics of the loss of CCAN protein levels and localization (Figure 2C).

**Figure 2:**
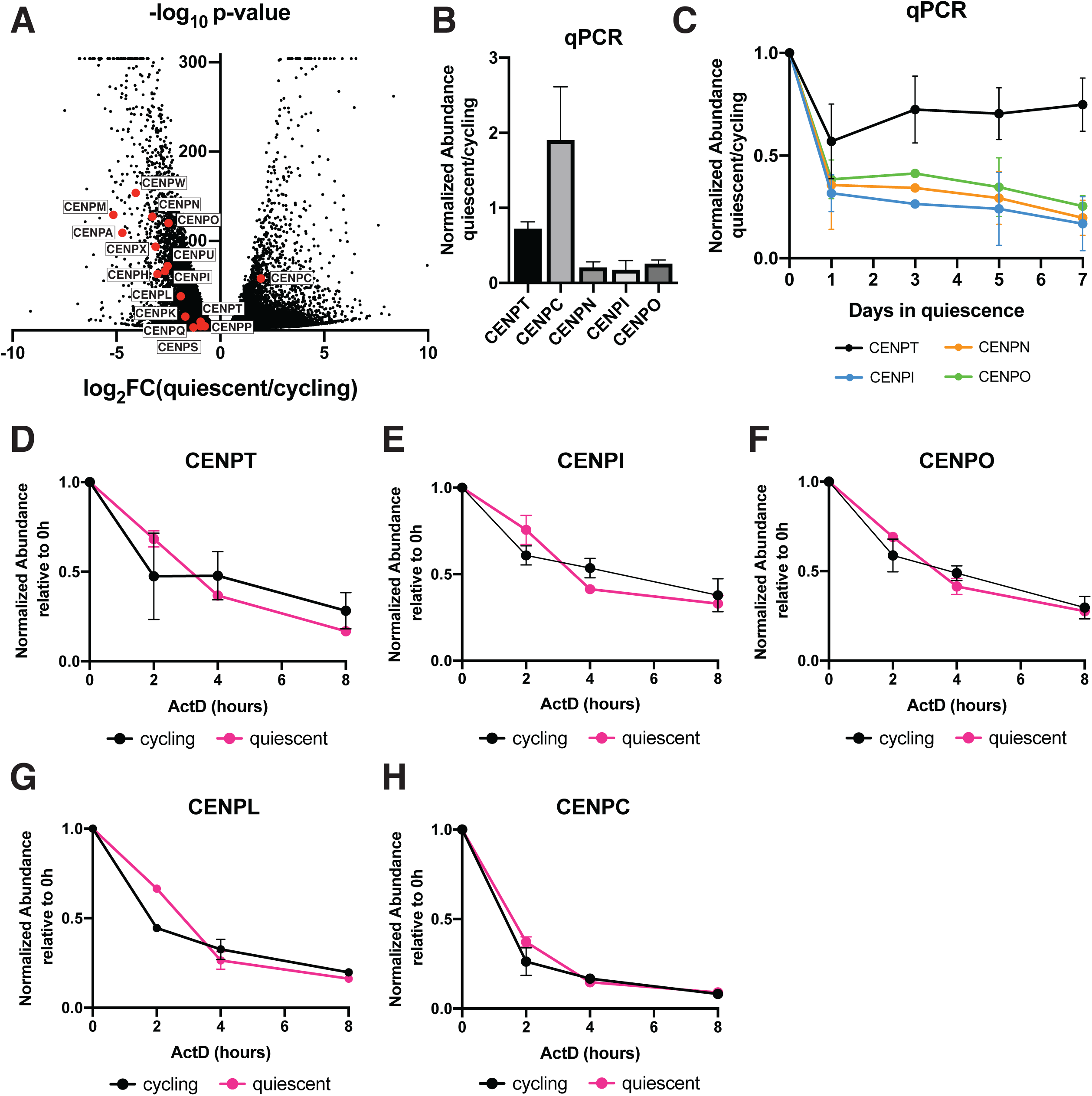
Quiescent cells regulate centromere behavior through a transcriptional program. A. Volcano plot comparing mRNA abundances in quiescent and cycling RPE1 cells as measured by RNA sequencing. Centromere components are highlighted in red. A p-value cut-off was imposed at p = 6.84E-305 for genes with p-values of 0. Genes with low read counts (total counts < 50) were excluded. B. Graph showing the fold change in mRNA abundance between cycling cells and cells in quiescence for 7 days for the indicated centromere component as quantified by qPCR. CT values were normalized to those of GAPDH before comparing quiescent and cycling values. Graph shows at least 3 biological replicates, with 3 technical replicates each for each centromere mRNA. Bars represent mean ± standard deviation. C. Graph showing mRNA abundance for centromere components over time as cells enter quiescence. Fold change is calculated for each quiescent time point by dividing by cycling value. CT values were normalized to those of GAPDH before comparing quiescent and cycling values. Graph shows at least 3 biological replicates, with 3 technical replicates each for each centromere mRNA. Bars represent mean ± standard deviation. D. Graph showing CENPT mRNA abundance over time after addition of 5µg/mL Actinomycin D in cycling and quiescent cells. Fold change is calculated for each ActD time point by dividing by untreated (0 hour) value for each respective condition (quiescent or cycling). CT values were normalized to those of GAPDH before comparing treated and untreated values. Graph shows 2 biological replicates, with 3 technical replicates each. Bars represent mean ± standard deviation. E. Graph showing CENPI mRNA abundance over time after addition of 5µg/mL Actinomycin D in cycling and quiescent cells. Fold change is calculated for each ActD time point by dividing by untreated (0 hour) value for each respective condition (quiescent or cycling). CT values were normalized to those of GAPDH before comparing treated and untreated values. Graph shows 2 biological replicates, with 3 technical replicates each. Bars represent mean ± standard deviation. F. Graph showing CENPO mRNA abundance over time after addition of 5µg/mL Actinomycin D in cycling and quiescent cells. Fold change is calculated for each ActD time point by dividing by untreated (0 hour) value for each respective condition (quiescent or cycling). CT values were normalized to those of GAPDH before comparing treated and untreated values. Graph shows 2 biological replicates, with 3 technical replicates each. Bars represent mean ± standard deviation. G. Graph showing CENPL mRNA abundance over time after addition of 5µg/mL Actinomycin D in cycling and quiescent cells. Fold change is calculated for each ActD time point by dividing by untreated (0 hour) value for each respective condition (quiescent or cycling). CT values were normalized to those of GAPDH before comparing treated and untreated values. Graph shows 2 biological replicates, with 3 technical replicates each. Bars represent mean ± standard deviation. H. Graph showing CENPC mRNA abundance over time after addition of 5µg/mL Actinomycin D in cycling and quiescent cells. Fold change is calculated for each ActD time point by dividing by untreated (0 hour) value for each respective condition (quiescent or cycling). CT values were normalized to those of GAPDH before comparing treated and untreated values. Graph shows 2 biological replicates, with 3 technical replicates each. Bars represent mean ± standard deviation.

Decreases in centromere mRNA levels could be due to downregulated transcription or to reduced RNA stability. To test whether CENP mRNAs were less stable in quiescent cells, we treated cycling and quiescent cells with the transcriptional inhibitor, actinomycin D, and measured the change in mRNA levels over time (Figure 2D-H). We found that centromere mRNAs showed similar rates of degradation in both cycling and quiescent cells. Thus, decreased centromere mRNA levels in quiescence are not due to decreased RNA stability, but instead reflects decreased transcription. Thus, the regulation of centromere disassembly during quiescence entry likely occurs, at least in part, through a rapidly regulated transcriptional program and the subsequent reduction in centromere protein levels.

### CENP-C retention supports centromere identity in quiescent cells

Because CENP-C was the only CCAN component to show continued transcription, protein abundance, and centromere localization in quiescent cells (Figure 1I, J, L, Figure 2A, B), we next considered whether its retention could play a role in maintaining centromere identity during quiescent arrest. We previously found that the continuous replenishment of the centromeric histone variant, CENP-A, during quiescence was required for chromosome segregation following reentry into the cell cycle^40^. As CENP-C is necessary for CENP-A deposition in cycling cells^48–51^, we tested whether depleting CENP-C would compromise CENP-A incorporation in quiescent cells. To do so, we depleted CENP-C using RNAi and monitored CENP-A levels by immunofluorescence using a HaloTag pulse-chase system^52^ (Figure S2C, D). This system relies on the conjugation of fluorescent small molecules to the HaloTag to distinguish between pre-existing and newly incorporated CENP-A. After treating quiescent cells with either CENP-C or control siRNAs for 24 hours, we blocked pre-existing Halo-CENP-A (Figure S2C). Cells were then allowed to remain in quiescence for 6 more days, during which newly made, unblocked HaloTag-CENP-A is deposited at centromeres (Figure S2C). We then compared total or newly deposited CENP-A between quiescent cells treated with CENP-C or control siRNAs by immunofluorescence. Although total centromeric CENP-A levels were unchanged (4% reduction; Figure S2E), the incorporation of new CENP-A was reduced upon loss of CENP-C (Figure S2F, G). CENP-C depletion led to a 15% reduction in CENP-A deposition over the course of 6 days (Figure S2F, G). As cells can remain quiescent for years, even this smaller reduction in CENP-A deposition in the absence of CENP-C could lead to the dramatic loss of CENP-A over longer periods of quiescent arrest and subsequent defects upon the resumption of cell division. Previous work has also suggested the presence of CENP-C-independent CENP-A deposition pathways that could function in parallel^53,54^. Thus, the lack of a complete abrogation of CENP-A deposition following CENP-C loss could reflect the use of such alternative pathways during quiescence or could also result from incomplete depletion of CENP-C in the first few days of siRNA treatment.

### *De novo* CENP-A deposition during quiescence exit occurs following the first mitosis

We next sought to analyze centromere dynamics during cell cycle reentry following quiescence exit. In quiescent cells, CENP-A is retained at the centromeres, but with reduced levels (Figure S1D, Figure 3B, C). We therefore considered whether full CENP-A levels would be regained upon quiescence release, and when the timing of new CENP-A deposition would occur. In cycling cells, new CENP-A deposition at centromeres is strictly regulated temporally, occurring once per cell cycle in early G1^49,55,56^. As quiescent cells enter G1 upon return to proliferation, we hypothesized that new CENP-A deposition could occur either immediately upon release at the first G1, non-canonically at a later stage in the cell cycle, or during the subsequent G1 following the first mitotic division. To test this, we monitored a GFP-tagged CENP-A cell line^57^ by live cell imaging after exit from quiescence (Figure 3A). Cells released from quiescence for 12 hours did not have significantly different levels of GFP-CENP-A at the centromeres relative to quiescent cells, with both showing around a 60 to 70 percent reduction in GFP centromere intensity relative to cycling cells (Figure 3B, C, D, E). These reduced levels persisted for an average of around 36 hours after release, until cells underwent their first mitosis (Figure 3D, E, F). Following completion of the first mitosis and entry into the subsequent G1, centromeric GFP-CENP-A levels increased dramatically (Figure 3D). Thus, cells released from quiescence undergo their first CENP-A deposition event during the G1 following the first mitotic division.

**Figure 3:**
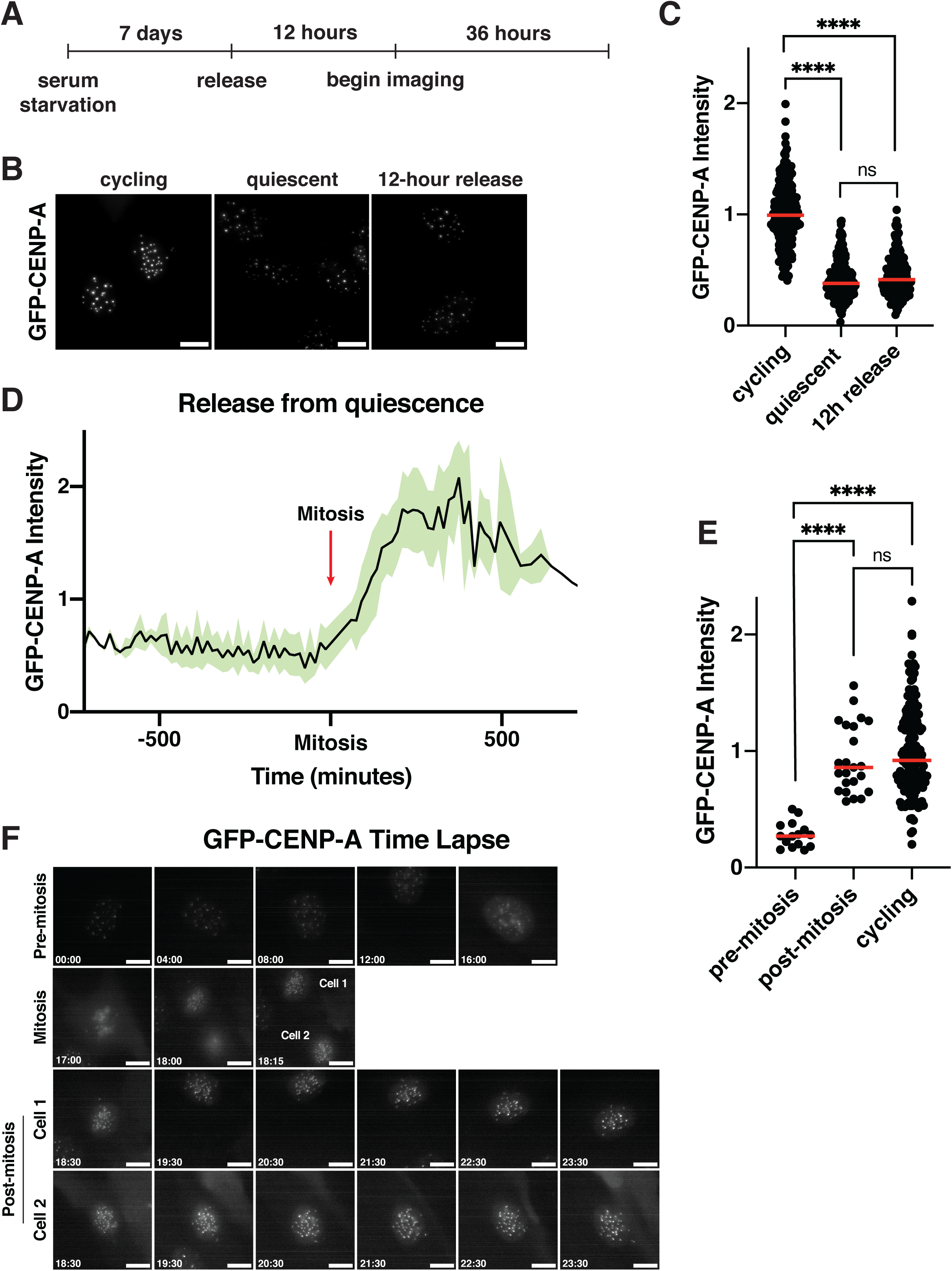
*De novo* CENP-A deposition during quiescence exit occurs following the first mitosis. A. Diagram showing experimental conditions for GFP-CENP-A quiescence exit live imaging experiments from Figure 3D, E, and F. B. Representative images of live GFP-CENP-A-expressing cells imaged for GFP from three different conditions: asynchronous cycling, 7 days of quiescence, or 12 hours after release from quiescence. Scale bar = 10 µm C. Graph quantifying cells from experiment in Figure 3B. GFP centromere intensity was measured in GFP-CENP-A-expressing cells from different conditions: asynchronous cycling, 7 days of quiescence, or 12 hours after release from quiescence. Cells were imaged live for GFP. Each point indicates the average centromere intensity level for 10 centromeres in a single cell, adjusted for background. Red line represents the median. Intensity values are normalized to cycling. p<0.0001 between cycling and quiescent and between cycling and 12h release, p = 0.1203 between quiescent and 12h release. **** represents p<0.0001, ns is not significant. n = 194, 196, 191 cells for cycling, quiescent, and 12 hour release, respectively D. Graph showing the normalized level of GFP-CENP-A intensity at the centromeres as cells exit quiescence. Red arrow indicates time of mitosis. X-axis spans from 12 hours before and 12 hours after mitosis. Black line shows the means and green shows mean ± standard deviation. Data is aggregated from 24 cells quantified over 15-minute timeframes from 2 separate biological replicates. E. Graph showing GFP-CENP-A intensity at the centromeres for cells released from quiescence before and after their first mitosis and for cycling cells (from Figure 3D). Pre-mitosis values are the average of all measurements from beginning of the live imaging or time of cell entry into imaging area to time of mitosis. Post-mitosis values are the average of all measurements from 1 hour after the end of mitosis to the end of the time course or exit of the cell from the imaging area. Cycling values are from asynchronous control cells from the last imaging time frame. Each point indicates the average centromere intensity level for 10 centromeres in a single cell, adjusted for background. Red line represents the median. Intensity values are normalized to cycling. p<0.0001 between pre-mitosis and post-mitosis and between pre-mitosis and cycling, p = 0.3236 between post-mitosis and cycling. **** represents p<0.0001, ns is not significant. F. Live imaging stills of a representative GFP-CENP-A cell exiting quiescence. Entry into mitosis is indicated. After mitosis, both cells are indicated and shown. Time units are hours:minutes.

As quiescent cells have reduced levels of CENP-A nucleosomes (Figure S1D, Figure 3B, C), we next considered whether cells would be able to restore the level of CENP-A typically observed at centromeres in cycling cells. We found that cells released from quiescence fully regained GFP-CENP-A levels equivalent to that of their cycling counterparts within a single deposition event, amounting to an approximately 3-fold increase of centromeric GFP-CENP-A (Figure 3E). Thus, cells exiting quiescence are able to regulate the amount of CENP-A deposited to restore homeostatic cycling levels of CENP-A within a single deposition cycle.

Together these results show that CENP-A levels are fully restored upon quiescence exit, but its redeposition does not occur in the first G1 following cell cycle reentry, but rather during the G1 following exit from the first mitotic division.

### Centromere reassembly during quiescence exit occurs in S phase but is independent of DNA replication

Unlike CENP-A, which persists at low levels in quiescent cells, most other CCAN components are completely lost during quiescent arrest, resulting in centromere disassembly (Figure 1A-H). As these CCAN proteins are required for chromosome segregation, they must be reassembled prior to the first mitosis. We therefore examined how the CCAN proteins are reassembled during the return to proliferation. In cells released from contact inhibition into full-serum media, CCAN component localization and protein levels were rapidly regained, starting within 24 hours of quiescence release (Figure S3A-F). To test at which stage of the cell cycle CCAN proteins are re-deposited at centromeres, we released cells from quiescence for 24 hours in the presence of 5-ethynyl-2’-deoxyuridine (EdU), a nucleotide analog that can be used to monitor progression through S phase via its incorporation during DNA replication. EdU positive cells, which have entered S phase, regained centromeric levels of CENP-T and CENP-O/P, whereas EdU negative cells did not (Figure 4A-F). In addition, the amount of EdU incorporation was closely correlated with CENP-T and CENP-O/P centromere intensities (Figure 4C, F). By contrast, CENP-C intensity was not correlated to EdU intensity and CENP-C levels increased in both EdU positive and EdU negative cells, although they showed a slightly greater relocalization in EdU positive cells (Figure S3G, H).

**Figure 4:**
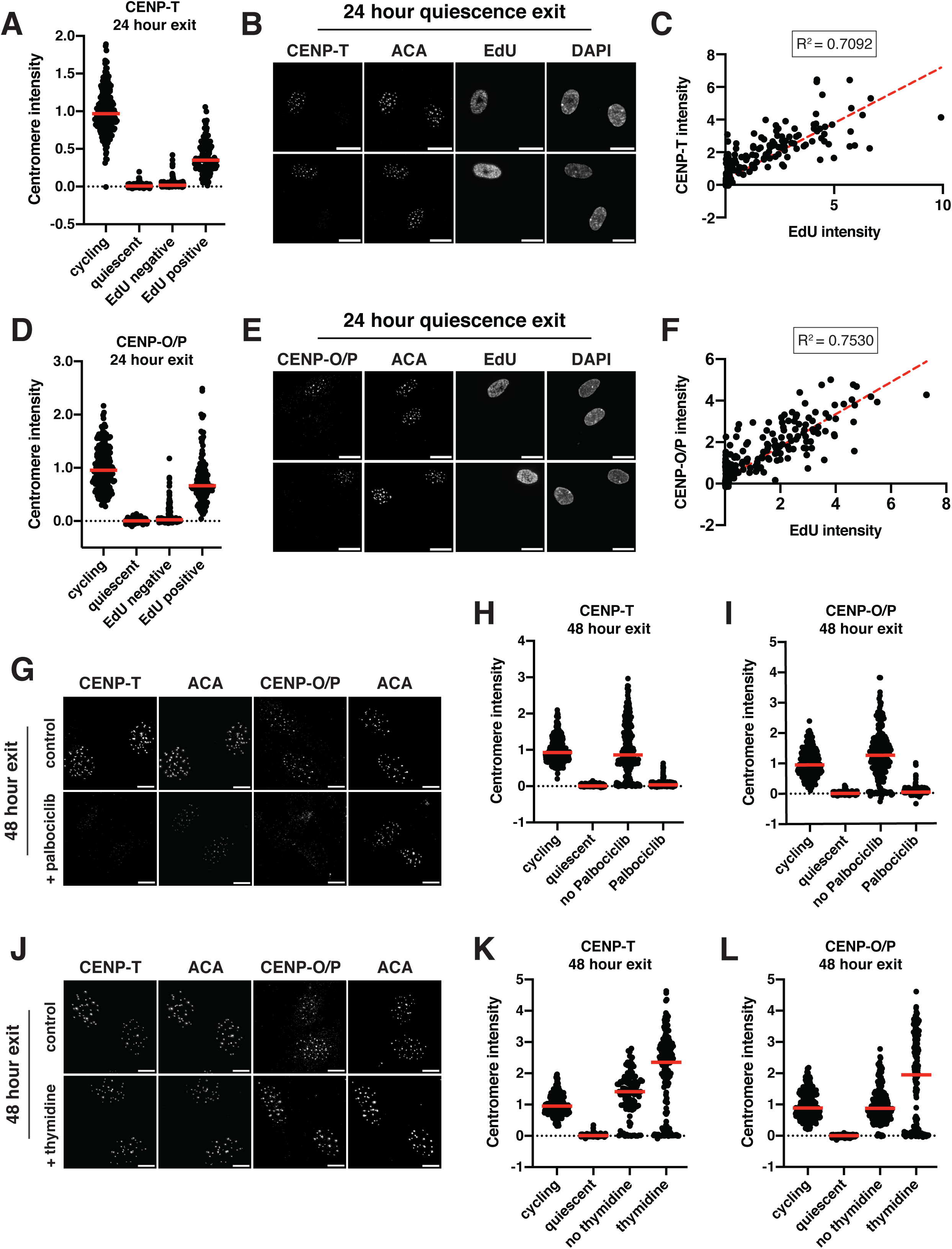
Centromere reassembly during quiescence exit occurs in S phase but is independent of DNA replication. A. Graph showing CENP-T centromere intensity levels for different conditions: cycling, 7 days of quiescence, and either EdU negative or EdU positive cells from cells released 24 hours from quiescence. Each point indicates the average centromere intensity level for all centromeres in a single cell, adjusted for background. Intensity values were normalized to cycling. Red line represents the median. Points were aggregated from 3 replicates. n = 202, 336, 202, and 107 cells for cycling, quiescent, 24 hours release EdU negative, and 24 hours release EdU positive respectively. All pairwise comparisons were significant with p < 0.0001 using unpaired t-test with Welch’s correction. B. Representative immunofluorescence images of cells released 24 hours from quiescence. EdU positive and negative cells are shown. Cells were stained with CENP-T and anti-centromere (ACA) antibodies. Scale bar = 20µm. C. Graph showing CENP-T centromere intensity plotted against EdU intensity for each cell. CENP-T intensity is the average CENP-T centromere intensity level for all centromeres in a cell, adjusted for background. EdU intensity is the mean EdU intensity in the nucleus of the same cell. Points were aggregated from 3 replicates. Values were normalized within each replicate. R^2^ = 0.7092. n = 309 cells. D. Graph showing CENP-O/P centromere intensity levels for different conditions: cycling, 7 days of quiescence, and either EdU negative or EdU positive cells from cells released 24 hours from quiescence. Each point indicates the average centromere intensity level for all centromeres in a single cell, adjusted for background. Intensity values were normalized to cycling. Red line represents the median. Points were aggregated from 3 replicates. n = 214, 389, 185, and 114 cells for cycling, quiescent, 24 hours release EdU negative, and 24 hours release EdU positive respectively. All pairwise comparisons were significant with p < 0.0001 using unpaired t test with Welch’s correction. E. Representative immunofluorescence images of cells released 24 hours from quiescence. EdU positive and negative cells are shown. Cells were stained with CENP-O/P and anti-centromere (ACA) antibodies. Scale bar = 20µm. F. Graph showing CENP-O/P centromere intensity plotted against EdU intensity for each cell. CENP-O/P intensity is the average CENP-O/P centromere intensity level for all centromeres in a cell, adjusted for background. EdU intensity is the mean EdU intensity in the nucleus of the same cell. Points were aggregated from 3 replicates. Values were normalized within each replicate. R^2^ = 0.753. n = 298 cells. G. Representative immunofluorescence images of cells released from quiescence for 48 hours with or without addition of 1 µM Palbociclib. Cells were stained for either CENP-T or CENP-O/P and anti-centromere (ACA) antibodies. Scale bar = 10 µm. H. Graph showing CENP-T centromere intensity for different conditions: cycling, quiescent, 48-hour release from quiescence without Palbociclib, and 48-hour release from quiescence with Palbociclib. Each point indicates the average centromere intensity level for all centromeres in a single cell, adjusted for background. Intensity values were normalized to cycling. Red line represents the median. Points were aggregated from 3 replicates. n = 353, 429, 237, and 280 for cycling, quiescent, no Palbociclib, and Palbociclib, respectively. There is no significant difference between cycling and no Palbociclib, p = 0.9691. All other pairwise comparisons were significant with p < 0.0001 using unpaired t test with Welch’s correction. I. Graph showing CENP-O/P centromere intensity for different conditions: cycling, quiescent, 48-hour release from quiescence without Palbociclib, and 48-hour release from quiescence with Palbociclib. Each point indicates the average centromere intensity level for all centromeres in a single cell, adjusted for background. Intensity values were normalized to cycling. Red line represents the median. Points were aggregated from 3 replicates. n = 244, 355, 215, and 256 for cycling, quiescent, no Palbociclib, and Palbociclib, respectively. All pairwise comparisons were significant with p < 0.0001 using unpaired t test with Welch’s correction. J. Representative immunofluorescence images of cells released from quiescence for 48 hours with or without addition of 2 mM thymidine. Cells were stained for either CENP-T or CENP-O/P and anti-centromere (ACA) antibodies. Scale bar = 10 µm. K. Graph showing CENP-T centromere intensity for different conditions: cycling, quiescent, 48-hour release from quiescence without thymidine, and 48-hour release from quiescence with thymidine. Each point indicates the average centromere intensity level for all centromeres in a single cell, adjusted for background. Intensity values were normalized to cycling. Red line represents the median. Points were aggregated from 3 replicates. n = 247, 343, 112, and 166 for cycling, quiescent, no thymidine, and thymidine, respectively. All pairwise comparisons were significant with p < 0.0001 using unpaired t test with Welch’s correction. L. Graph showing CENP-O/P centromere intensity for different conditions: cycling, quiescent, 48-hour release from quiescence without thymidine, and 48-hour release from quiescence with thymidine. Each point indicates the average centromere intensity level for all centromeres in a single cell, adjusted for background. Intensity values were normalized to cycling. Red line represents the median. Points were aggregated from 3 replicates. n = 260, 328, 149, and 190 for cycling, quiescent, no thymidine, and thymidine, respectively. There is no significant difference between cycling and no thymidine, p = 0.9002. All other pairwise comparisons were significant with p < 0.0001 using unpaired t test with Welch’s correction.

Given the correlation between centromere reassembly and EdU incorporation, we next evaluated whether CCAN deposition upon quiescence exit requires entry into S phase. To test this, we released quiescent cells into the cell cycle for 48 hours in the presence of Palbociclib, a CDK4/6 inhibitor that prevents S phase entry and induces a G1 arrest. Cells treated with Palbociclib did not enter S phase and no longer incorporated EdU (Figure S3I, J). Accordingly, Palbociclib treatment prevented the redeposition of CENP-T and CENP-O/P, whereas CENP-C relocalization was unaffected (Figure 4G, H, I; Figure S3K). Thus, preventing S phase entry upon quiescence exit prevents centromere reassembly.

To test whether centromere reassembly requires DNA replication, we next released cells from quiescence for 48 hours in the presence or absence of thymidine, which prevents DNA replication and arrests cells in early S phase^58,59^. Cells released from quiescence in the presence of thymidine were able to re-gain CENP-T, CENP-O/P, and CENP-C protein localization (Figure 4J, K, L; Figure S3L). Thus, centromere deposition does not require DNA replication. In addition, this experiment confirms that CCAN deposition indeed occurs during S phase and does not require entry into a later cell cycle stage.

Interestingly, although untreated cells released from quiescence recovered normal cycling levels of CCAN at the centromeres, we observed even higher centromeric CCAN protein intensity in thymidine-treated cells relative to untreated cells (Figure 4K, L). This suggests that cells can usually restore homeostatic levels of CENP-T and CENP-O/P upon return to growth, but that extending the time spent in S phase can lead to additional centromere protein deposition. Indeed, prolonging S phase even in normal cycling cells through 48-hour thymidine incubation also resulted in increased centromere protein deposition (Figure S3M, N). Thus, cells regulate the appropriate level of centromere protein deposition under normal circumstances, but prolonging the stage at which deposition occurs and access to potential deposition factors can disturb this regulatory control.

Together, these results show that centromere reassembly following exit from quiescence occurs in S phase and that this deposition is independent of DNA replication.

## Discussion

Quiescence is a unique cellular state that requires a cell to enforce a prolonged arrest while maintaining its capacity to divide again in the future. Here, we show how cells modify a key cellular structure, the centromere, to meet the demands of the quiescent state. The centromere is crucial for chromosome segregation and cell division in cycling cells. In quiescent cells, which no longer divide and do not require centromere function, we find that the centromere is disassembled through the complete loss of specific centromere components, such as CENP-T, CENP-K, CENP-L, and CENP-O/P. This loss is regulated in part by a rapidly enacted transcriptional program, which downregulates centromere component transcription within 24 hours of serum starvation. However, quiescent cells must also preserve the centromere to be able to segregate their chromosomes upon return to the cell cycle. We show that selected centromere components required for maintaining centromere identity, such as CENP-A and CENP-C, are retained. Thus, the centromere is a prime example of how cells modify and regulate a cellular structure to enable the requirements of quiescence.

Prior work on cell division components has focused primarily on cycling cells. However, understanding the logic of cell division in an intact organism requires a consideration of the diverse physiological circumstances in which cells exist, including non-dividing states. Our work highlights how cellular quiescence can be used to obtain novel insights into centromere biology. In proliferating cells, components of the CCAN are constitutively localized to the centromere in all stages of the cell cycle. In contrast, we show that quiescence provides a unique context in which the centromere is largely disassembled. By analyzing the dynamics and requirements of how this disassembly occurs, we observed distinct behaviors amongst the CCAN components. CENP-T, CENP-L, CENP-K, and CENP-O/P, which are usually constitutively present at centromeres, are completely lost within days of quiescence induction. Conversely, CENP-A and CENP-C are retained as a minimal unit required to fully restore centromere protein localization upon the return to growth.

Release from quiescence and return to growth also provides a unique opportunity to study centromere reassembly and centromere protein deposition. Whereas previous work studied centromere protein deposition in the context of cycling cells^60^, where pre-existing pools of protein can be difficult to distinguish from newly deposited protein, quiescence exit allows for the observation of completely *de novo* centromere protein deposition. We find that new CENP-T and CENP-O/P are deposited at S phase, independent of DNA replication, whereas increases in CENP-C levels can occur earlier. In addition, we show here that *de novo* CENP-A deposition in cells that exit quiescence occurs in the G1 following the first mitosis, and not in the G1 immediately following quiescence release.

In addition to defining the timing of centromere protein deposition, the return to growth from quiescence also provides insight on how cells regulate the homeostatic levels of centromere proteins at centromeres and the stoichiometry of deposition. Whereas cycling cells are expected to halve CENP-A levels during DNA replication and double them again during G1, such that each pre-existing CENP-A would recruit a single new molecule of CENP-A, release from quiescence provides a unique situation in which more CENP-A must be recruited to restore homeostatic levels. Indeed, we observe an around 3-fold increase in CENP-A at the centromere during the first deposition event following quiescence exit and cells exiting quiescence regain cycling-level centromeric CENP-A within only one cell division. These results highlight an interesting aspect of the stoichiometry of CENP-A deposition, where deposition of new CENP-A need not always occur at a one-to-one ratio. Similarly, the levels of other CCAN proteins, CENP-T and CENP-O/P, also appear to be closely regulated during return to growth. Cells released from quiescence regulate the transition from the complete absence of these CCAN proteins to achieve cycling levels. However, if the deposition period is lengthened by arresting cells in S phase, this regulation is disrupted and CCAN protein levels can overtake normal cycling amounts. This suggests that the amount of CCAN protein at the centromeres is in part regulated by the length of time spent in a state permissive to deposition.

In all, this work not only highlights important paradigms in the regulation of cellular structures during quiescence, but also uncovers key aspects of centromere biology within a novel cellular context.

## Acknowledgments

We thank members of the Cheeseman lab for feedback throughout the process. We would also like to thank members of the Whitehead Genome Technology Core. This work was supported by a grant to I.M.C. from the NIH/National Institute of General Medical Sciences (R35GM126930).

## Author Contributions

Conceptualization: OM, IMC; Methodology, Investigation, Validation: OM did experiments with help from KCS for HaloTag CENP-A experiments, BM for qPCR time course replicates, and NT for additional experiments; Supervision: IMC; Funding acquisition: IMC; Writing and editing: OM, IMC.

## Method Details

### Cell Lines

hTERT RPE-1 and NIH-3T3 cell lines were cultured in Dulbecco’s modified Eagle medium (DMEM) supplemented with either 100 U/ml penicillin and streptomycin and 2mM L-glutamine with (growth) or without (serum starvation) 10% fetal bovine serum (FBS). Cells were grown at 37°C with 5% CO_2_. hTERT RPE-1 cells are hTERT-immortalized retinal pigment epithelial cells of female origin. The NIH-3T3 cell line is derived from a male mouse embryo. Cells were tested regularly for mycoplasma contamination.

### Cell culture and reagents

For quiescence induction, cells were allowed to grow until reaching confluence. Cells were then left to contact inhibit for 1-2 days before serum starvation. For serum starvation, full serum media was removed from cells, which were then washed once with PBS (phosphate buffered saline), before replacing with serum-free DMEM. Media was then changed every 24 hours until cells were harvested. Quiescent cells were serum starved for 7 days unless otherwise noted. For quiescence release, quiescent cells were trypsinized or treated with PBS + 5mM EDTA before replating at lower density in full serum (10% FBS) media for the indicated amounts of time.

Drugs used on cell lines were: 5-ethynyl-2’-deoxyuridine (EdU; 10 µM; Vector Laboratories), actinomycin D (5 µg/mL; Santa Cruz Biotechnology), thymidine (2 mM; Sigma-Aldrich), Palbociclib (1 µM; Selleck), Janelia Fluor HaloTag Ligand (125 nM or 30 nM; Promega).

### Immunofluorescence

Cells were fixed with either 4% formaldehyde diluted in phosphate-buffered saline (PBS) + 0.5% TritonX-100 or in cold Methanol. After washing with PBS + 0.1% TritonX-100 and incubation in blocking buffer (20 mM Tris-HCl, 150 mM NaCl, 0.1% Triton X-100, 3% bovine serum albumin, 0.1% NaN_3_, pH 7.5) for 30 minutes or overnight, primary antibody was added for 1 hour or overnight. Cells were then washed before addition of Cy2-, Cy3-, or Cy5-conjugated secondary antibodies (Jackson ImmunoResearch Laboratories) for 1 hour, followed by 10 minutes of 1 µg/ml Hoechst-33342 (Sigma-Aldrich) diluted in 0.1% PBS-Tx to stain DNA. For cells with EdU staining, after incubation with secondary antibody, Click-iT buffer (100 mM Tris pH 8.0, 1 mM CuSO_4_, 5 µM AlexaFluor azide (Life Technologies), 100 mM Ascorbic Acid) was added for 30 minutes before washing and Hoechst staining. Coverslips were mounted onto slides with ProLong^TM^ Gold antifade reagent (Invitrogen). Microscopy was conducted using DeltaVision Ultra High-Resolution microscope and images were analyzed with Fiji (ImageJ, NIH) and CellProfiler^61^. Images were deconvolved on the DeltaVision. A list of antibodies and their dilutions can be found in Supplementary Table 1^43,62,63^.

### Live-cell Imaging

For live-cell imaging, GFP-CENP-A-tagged cells were induced to enter quiescence by contact inhibition and 7 days of serum starvation. Quiescent cells were then released from quiescence and replated onto 12-well glass-bottom plates (Cellvis, P12-1.5P) and centrifuged to promote adherence. 1 hour prior to imaging, the media was changed to CO_2_-independent media (Gibco) supplemented with 10% FBS, 100 U/ml penicillin and streptomycin, and 2 mM L-glutamine. Cells were imaged 12 hours after replating, for 36 hours with 15-minute time points. Imaging for GFP was conducted at 37°C on a Nikon eclipse microscope (40X). Images were analyzed with Fiji (ImageJ, NIH).

### Determining EdU positive and negative cells

Maximally projected images were processed by CellProfiler, where the EdU channel was measured within nuclear boundaries for each cell. To find the threshold EdU value to determine positive and negative cells, mean nuclear EdU intensities for all cells were used to fit a Gaussian Mixture Model (GMM) with 2 components. GMM was then used to determine threshold based on predictions from a range of values. Code was developed using help from ChatGPT and is available on Github.

### Halo-Tag Pulse-chase

HaloTag-CENP-A cells were induced to enter quiescence by contact inhibition and serum starvation. 4 days after start of serum starvation, cells were treated with control or CENP-C siRNAs. 24 hours later, pre-existing CENP-A was blocked by incubation with 125 nM of Halo JF549. After 2 days, siRNAs were reapplied. After 4 more days of quiescence (11 total days of quiescence), cells were fixed. Immunofluorescence was conducted as above, however during incubation with secondary, cells were also incubated with 30 nM Halo JF646 for one hour before washing and Hoechst staining.

### Western Blot

Cells were lysed with urea lysis buffer (50 mM Tris pH 7.5, 150 mM NaCl, 0.5% NP-40, 0.1% sodium dodecyl sulfate (SDS), 6.5 M Urea, 1X Complete EDTA-free protease inhibitor cocktail (Roche), 1 mM phenylmethylsulfonyl fluoride (PMSF)) on ice for 25 minutes, followed by centrifugation to remove cellular debris. Then, Laemmli sample buffer was added with 2-mercaptoethanol and samples were heated at 95°C for 5 minutes. Samples were then loaded onto acrylamide gels for SDS-PAGE, followed by 1 hour or overnight transfer onto nitrocellulose membranes. Membranes were incubated for 1 hour in blocking buffer (2.5% milk in Tris-Buffered Saline (TBS) + 0.1% Tween-20), then 1 hour or overnight in primary antibodies, then 1 hour in either HRP-conjugated secondary antibodies (GE Healthcare; Digital) or in IRDye secondary antibodies (LI-COR). For imaging, blots were either incubated in clarity-enhanced chemiluminescence substrate (Bio-Rad) and imaged using the KwikQuant Imager (Kindle Biosciences) or directly imaged using the Odyssey CLx Imager (LI-COR). β-actin is used as a loading control. Antibodies are listed in Supplementary Table 1.

### Flow Cytometry Analyses

For DNA content analysis, cells were harvested, resuspended in 1 ml of PBS, and fixed with addition of 9 ml of ice-cold ethanol. Cells were then resuspended in PBS to rehydrate, followed by incubation in blocking buffer (20 mM Tris-HCl, 150 mM NaCl, 0.1% Triton X-100, 3% bovine serum albumin, 0.1% NaN_3_, pH 7.5) for 30 minutes. After blocking, cells were resuspended in PBS containing 10 µg/ml RNase A and 20 µg/ml propidium iodide (PI, Invitrogen) and incubated for 30 minutes before analysis by flow cytometry. PI signal was measured on an LSRFortessa (BD Biosciences) flow cytometer. Results were analyzed with FlowJo software. Data was collected on at least 10,000 cells for each condition per experiment.

### siRNA treatment

Custom siRNAs against CENP-C (5’-GAACAGAAUCCAUCACAAAUU), and a non-targeting control pool (D-001206-13) were obtained from Dharmacon. siRNAs were used at a final concentration of 50 nM. siRNAs and Lipofectamine RNAiMAX (Invitrogen) were mixed at equal volume in Opti-MEM Reduced Serum Medium (ThermoFisher), vortexed, and allowed to incubate for 20 minutes before adding to cells. Media was changed between 24 hours later back to no serum media for quiescence.

### RNA isolation, Reverse Transcription, qPCR, and RNA sequencing

Cells were lysed in 400 µl of TRIzol RNA isolation reagent (ThermoFisher) and frozen at −80°C or used directly for the next step. 120 µl of chloroform was added to samples and tubes were vortexed vigorously for at least 30 seconds. After centrifuging at 4°C, the aqueous phase was mixed with an equal volume of chloroform. Samples were vortexed vigorously, centrifuged at 4°C, and aqueous phase was again transferred to a new tube. GlycoBlue Coprecipitant (Invitrogen), 5 M NaCl, and equal volume of isopropanol were added, and samples were incubated on dry ice for 30 minutes. After centrifugation, pellets were isolated, washed with 75% Ethanol and resuspended in water. All reagents were RNase free. Maxima First Strand cDNA Synthesis Kit for RT-qPCR (Thermo Scientific) was used for reverse transcription according to manufacturer instructions. For qPCR, cDNAs were diluted and mixed with 1 µM primers and 2X SYBR Green PCR Master Mix (Thermo Fischer Scientific) in 384-well plates. Three technical replicates per cDNA sample and primer pair were used. Primer sequences can be found in Supplementary Table 1.

For RNA sequencing: rRNA depletion and library preparation were done with the KAPA RNA HyperPrep Kit with RiboErase (HMR) (Roche KK8560) according to manufacturer instructions. Sequencing was performed on an Illumina NovaSeq SP. For analysis, reads were aligned to the human (gencode v25, GRCh38) genome using hisat2. Expression matrix was generated using featureCounts. Genes with low read count (less than 50) were filtered out. Differential expression analysis was conducted with DESeq2.

### Quantification and Statistical Analysis

Quantification of fluorescence intensity for immunofluorescence experiments was conducted on unprocessed, maximally projected images using FIJI/Image J or using a custom CellProfiler and Python pipeline. These images were acquired using the same microscope and acquisition settings on the same day, unless otherwise noted in the figure legends. Statistical analyses were performed using Prism (GraphPad Software). Details of statistical tests and sample sizes are provided in the figure legends.

---

**Supplementary Figure 1:**
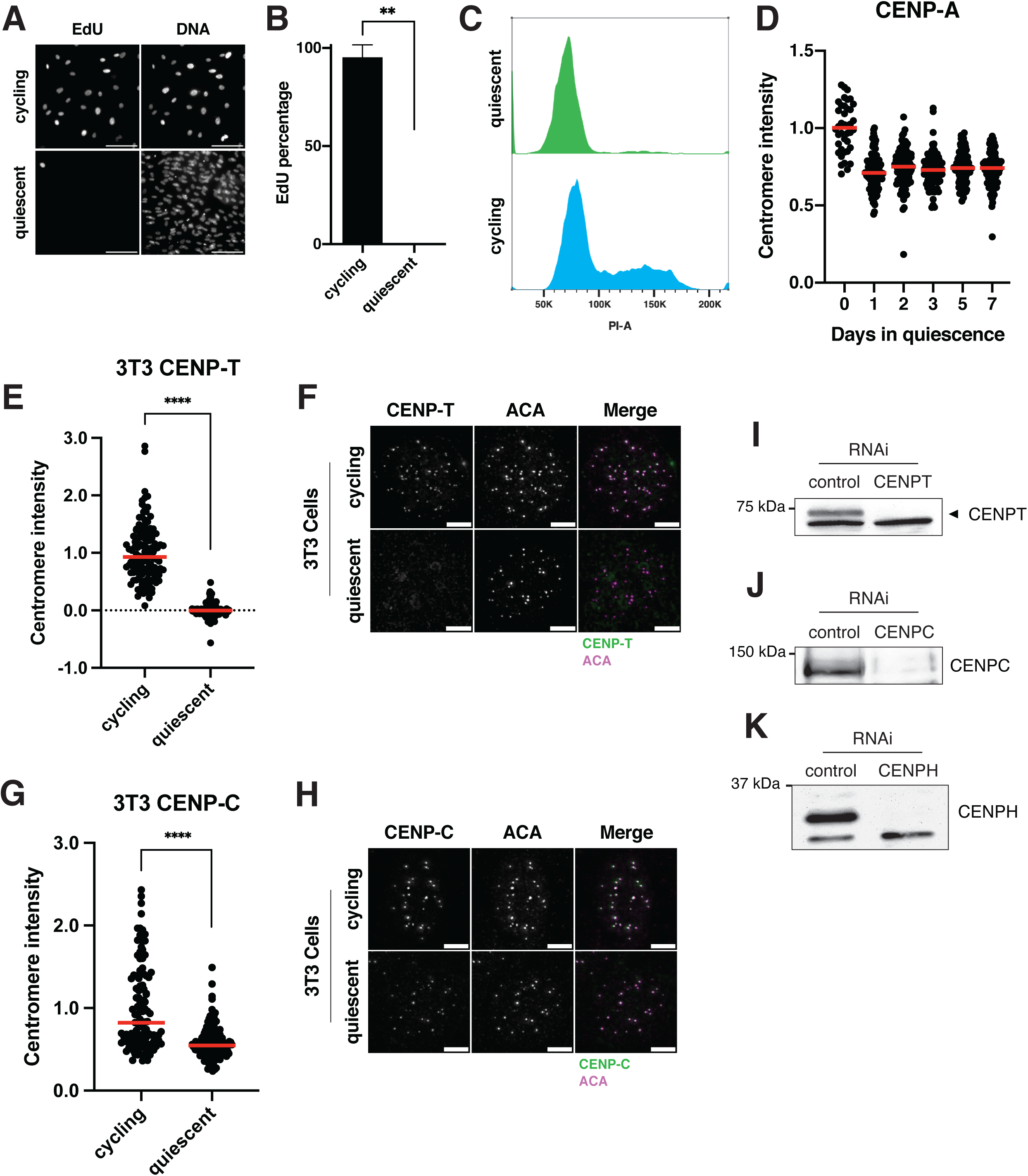
Controls for quiescence induction and conservation of quiescent centromere behavior in mouse cells. A. Representative images of EdU staining of cycling and quiescent cells. Cells were incubated for 48 hours in 5-ethynyl-2’ deoxyuridine (EdU), a nucleotide analog that monitors DNA replication and progression through the cell cycle. Scale bar = 100µm. B. Graph showing the percentage of EdU positive cells for the indicated condition. Cells were incubated for 48 hours in EdU. Bars represent mean ± standard deviation of three replicates. Mean of cycling is 95.12, mean of quiescence is 0.2. ** represents p = 0.0015. C. Histogram showing the distribution of propidium iodide (PI) staining for cycling cells (blue) or cells in quiescence for 7 days (green) as measured by flow cytometry. D. Graph showing CENP-A centromere intensity level over time of quiescence entry. Each point indicates the average centromere intensity level for all centromeres of a single cell, adjusted for background. Intensity values were normalized to day 0. Red line represents the median. Points were from 1 replicate, as this result has already been previously shown. n = 41, 110, 94, 84, 110, and 93 cells for 0, 1, 2, 3, 5, and 7-day time points respectively. E. Graph showing CENP-T centromere intensity levels in cycling and quiescent mouse 3T3 cells. Each point indicates the average centromere intensity level for a single cell, adjusted for background. Intensity values were normalized to cycling condition. Red line represents the median. Points were aggregated from 2 replicates. n = 121 and 123 cells for cycling and quiescent respectively. **** represents p<0.0001. F. Representative immunofluorescence images of cycling and quiescent mouse 3T3 cells. Cells were stained with mouse CENP-T and anti-centromere (ACA) antibodies. Scale bar = 5µm. G. Graph showing CENP-C centromere intensity levels in cycling and quiescent mouse 3T3 cells. Each point indicates the average centromere intensity level for a single cell, adjusted for background. Intensity values were normalized to cycling condition. Red line represents the median. Points were aggregated from 2 replicates. n = 121 and 136 cells for cycling and quiescent respectively. **** represents p<0.0001. H. Representative immunofluorescence images of cycling and quiescent mouse 3T3 cells. Cells were stained with mouse CENP-C and anti-centromere (ACA) antibodies. Scale bar = 5µm. I. Western blot of cells treated with 50 nM CENPT or control siRNAs. Blot shows the banding pattern for CENP-T antibody. CENP-T is the upper band. Blot was incubated in CENP-T antibody. J. Western blot of cells treated with 50 nM CENPC or control siRNAs. Blot shows the banding pattern for CENP-C antibody. Blot was incubated in CENP-C antibody. K. Western blot of cells treated with 50 nM CENPH or control siRNAs. Blot shows the banding pattern for CENP-H antibody. CENP-H is the upper band. Blot was incubated in CENP-H antibody.

**Supplemental Figure 2:**
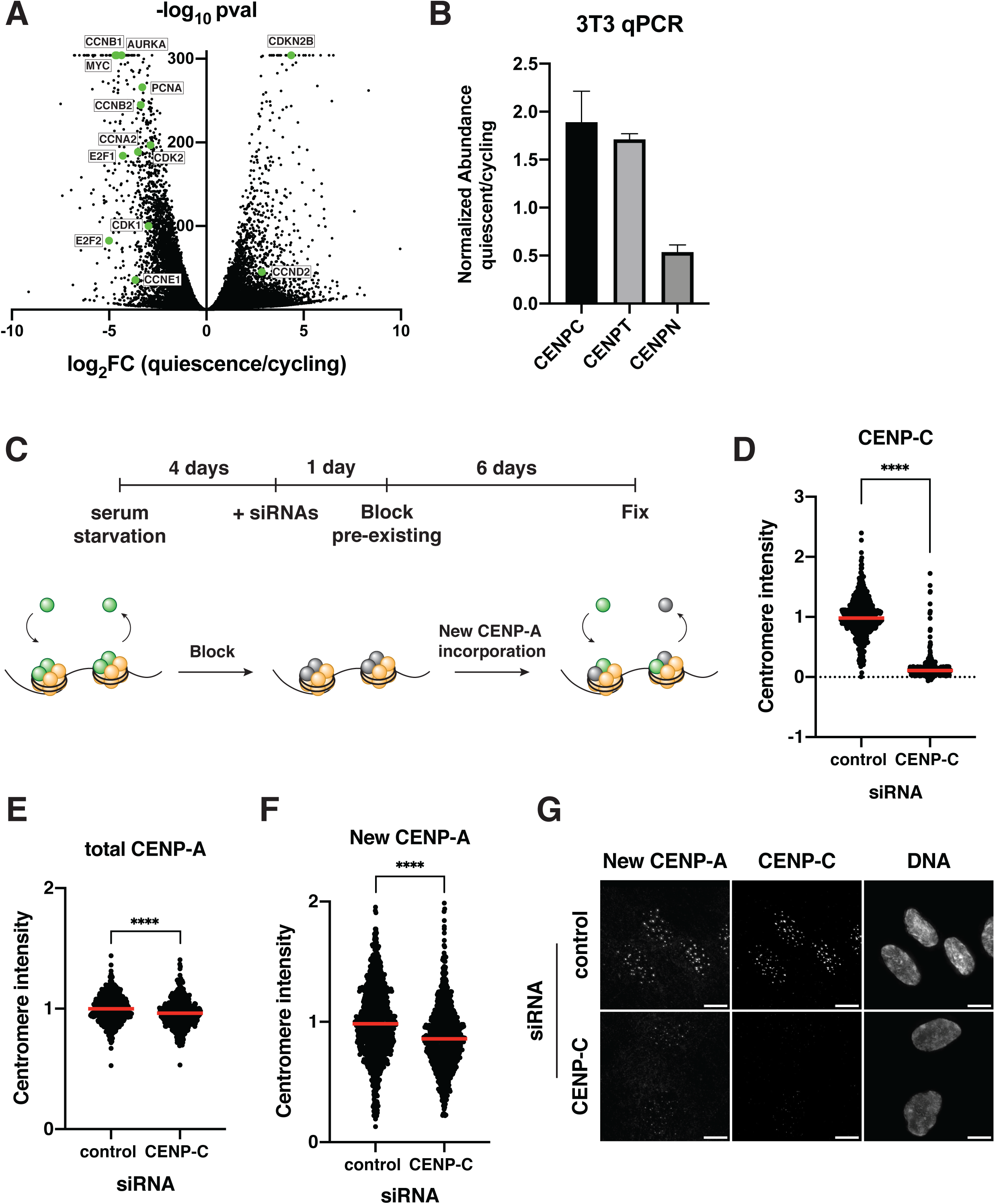
CENP-C contributes to CENP-A deposition in quiescent cells. A. Volcano plot comparing mRNA abundances in quiescent and cycling RPE1 cells as measured by RNA sequencing. Positive controls, including certain cyclins, cyclin-dependent kinases, and other proliferation factors, are highlighted in green. A p-value cut-off was imposed at p = 6.84E-305 for genes with p-values of 0. Genes with low read counts (total counts < 50) were excluded. B. Graph showing the fold change in mRNA abundance in mouse 3T3 cells between cycling cells and cells in quiescence for 7 days for the indicated centromere component as quantified by qPCR. CT values were normalized to those of GAPDH before comparing quiescent and cycling values. Graph shows at least 3 biological replicates, with 3 technical replicates each for each centromere mRNA. Bars represent mean ± standard deviation. C. Schematic showing experimental design for HaloTag pulse-chase experiments. Unblocked CENP-A is shown in green, blocked CENP-A in gray and other histones in yellow. More experimental details can be found in the methods section. D. Graph showing CENP-C intensity at the centromeres after 7 days of RNAi treatment. Cells were fixed and stained with CENP-C antibody at the end of the experiment from S2C. Each point indicates the average centromere intensity level for all centromeres in a single cell, adjusted for background. Intensity values were normalized to control. Points were aggregated from 3 replicates. **** indicates p<0.0001. n = 486, 432 for control and CENP-C RNAi conditions respectively E. Graph showing total CENP-A intensity at the centromeres after 7 days of RNAi treatment. Cells were fixed and stained with CENP-A antibody at the end of the experiment from S2C. Each point indicates the average centromere intensity level for all centromeres in a single cell, adjusted for background. Intensity values were normalized to control. Points were aggregated from 3 replicates. **** indicates p<0.0001. n = 481, 447 for control and CENP-C RNAi conditions respectively. F. Graph showing new, unblocked CENP-A intensity at the centromeres 6 days after blocking pre-existing CENP-A and after 7 days of RNAi treatment. Cells were fixed and stained JF646 at the end of the experiment from S2C. Each point indicates the average centromere intensity level for all centromeres in a single cell, adjusted for background. Intensity values were normalized to control. Points were aggregated from 3 replicates. **** indicates p<0.0001. n = 966, 879 for control and CENP-C RNAi conditions respectively. G. Representative immunofluorescent images showing levels of new unblocked CENP-A and CENP-C at the end of the pulse-chase experiment described in S2C. Cells were treated with either control or CENP-C siRNA. Scale bar = 10 µm.

**Supplementary Figure 3:**
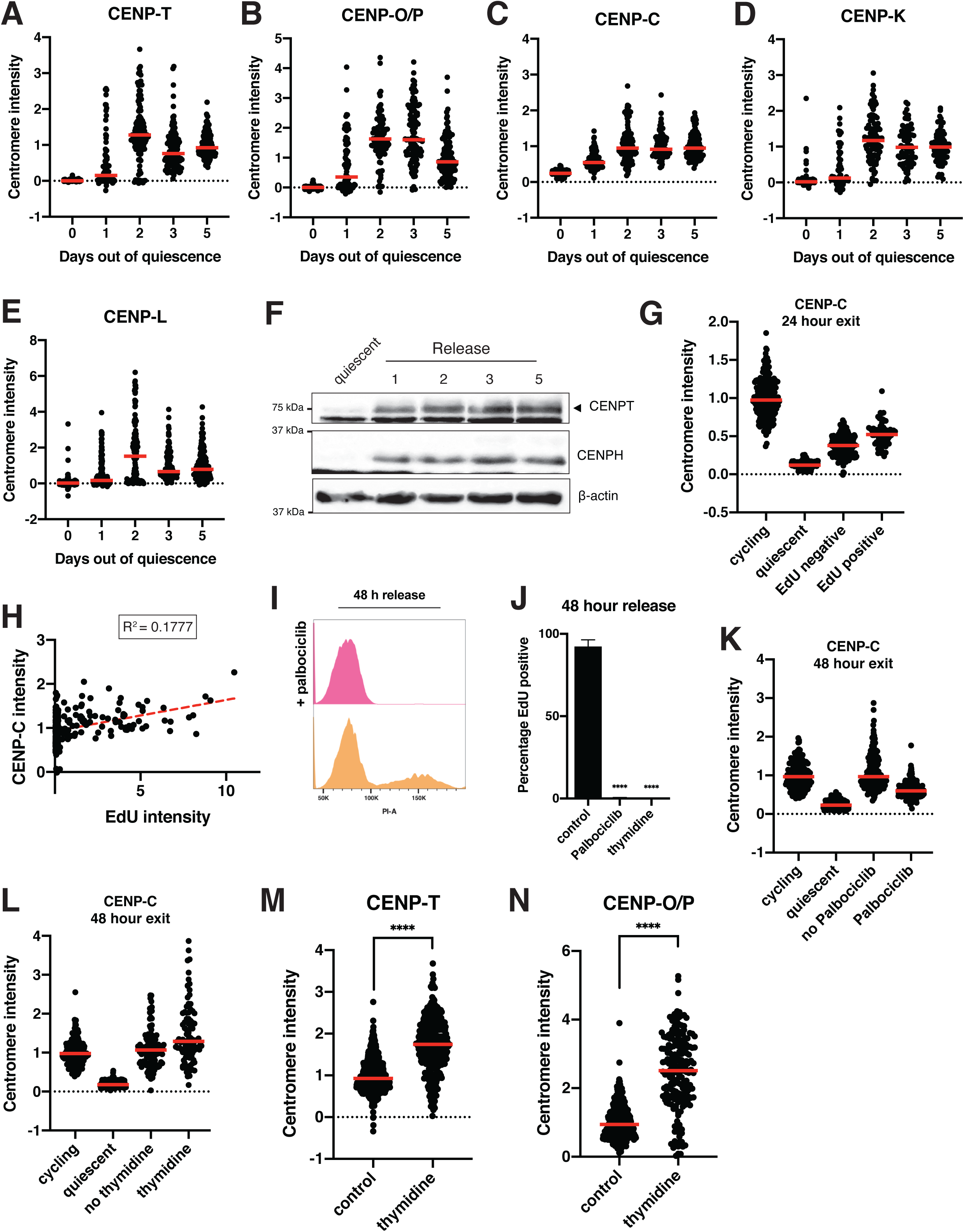
The centromere is rapidly reassembled upon cell cycle reentry. A. Graph showing CENP-T centromere intensity level over time of quiescence exit. Each point indicates the average centromere intensity level for all centromeres in a single cell, adjusted for background. Intensity values were normalized to day 5. Red line represents the median. Points were aggregated from 2 replicates. n = 232, 94, 116, 115, and 138 cells for 0, 1, 2, 3, and 5-day time points respectively. B. Graph showing CENP-O/P centromere intensity level over time of quiescence exit. Each point indicates the average centromere intensity level for all centromeres in a single cell, adjusted for background. Intensity values were normalized to day 5. Red line represents the median. Points were aggregated from 2 replicates. n = 238, 90, 94, 111, and 110 cells for 0, 1, 2, 3, and 5-day time points respectively. C. Graph showing CENP-C centromere intensity level over time of quiescence exit. Each point indicates the average centromere intensity level for all centromeres in a single cell, adjusted for background. Intensity values were normalized to day 5. Red line represents the median. Points were aggregated from 2 replicates. n = 180, 122, 133, 100, and 144 cells for 0, 1, 2, 3, and 5-day time points respectively. D. Graph showing CENP-K centromere intensity level over time of quiescence exit. Each point indicates the average centromere intensity level for all centromeres in a single cell, adjusted for background. Intensity values were normalized to day 5. Red line represents the median. Points were aggregated from 2 replicates. n = 179, 67, 89, 82, and 80 cells for 0, 1, 2, 3, and 5-day time points respectively. E. Graph showing CENP-L centromere intensity level over time of quiescence exit. Each point indicates the average centromere intensity level for all centromeres in a single cell, adjusted for background. Intensity values were normalized to day 5. Red line represents the median. Points were aggregated from 2 replicates. n = 203, 177, 132, 135, and 182 cells for 0, 1, 2, 3, and 5-day time points respectively. F. Western blots of cells release from quiescence for the indicated amount of days. Blots were incubated with CENP-T and CENP-H antibodies. β-actin is used as a loading control. G. Graph showing CENP-C centromere intensity levels for different conditions: cycling, 7 days of quiescence, and either EdU negative or EdU positive cells from cells released 24 hours from quiescence. Each point indicates the average centromere intensity level for all centromeres in a single cell, adjusted for background. Intensity values were normalized to cycling. Red line represents the median. Points were aggregated from 3 replicates. n = 199, 300, 206, and 66 cells for cycling, quiescent, 24 hours release EdU negative, and 24 hours release EdU positive respectively. All pairwise comparisons were significant with p < 0.0001 using unpaired t-test with Welch’s correction. H. Graph showing CENP-C centromere intensity plotted against EdU intensity for each cell. CENP-C intensity is the average CENP-C centromere intensity level for all centromeres in a cell, adjusted for background. EdU intensity is the mean EdU intensity in the nucleus of the same cell. Points were aggregated from 3 replicates. Values were normalized within each replicate. R^2^ = 0.1777. n = 272 cells. I. Histogram showing the distribution of propidium iodide (PI) staining for cells release 48 hours from quiescence without (orange) and with (pink) addition of Palbociclib, as measured by flow cytometry J. Graph showing the percentage of EdU positive cells for the indicated conditions. Cells were released from quiescence into full serum media and incubated for 48 hours in EdU with or without Palbociclib or thymidine. Bars represent mean ± standard deviation of 4 replicates for control, 3 replicates for thymidine, and 2 replicates for Palbociclib. Mean of control is 92.03, mean of Palbociclib is 0.3776, mean of thymidine is 0. **** represents p < 0.0001. K. Graph showing CENP-C centromere intensity for different conditions: cycling, quiescent, 48-hour release from quiescence without Palbociclib, and 48-hour release from quiescence with Palbociclib. Each point indicates the average centromere intensity level for all centromeres in a single cell, adjusted for background. Intensity values were normalized to cycling. Red line represents the median. Points were aggregated from 2 replicates. n = 174, 227, 198, and 207 for cycling, quiescent, no Palbociclib, and Palbociclib, respectively. There is no significant difference between cycling and no Palbociclib, p = 0.056. All other pairwise comparisons were significant with p < 0.0001 using unpaired t test with Welch’s correction. L. Graph showing CENP-C centromere intensity for different conditions: cycling, quiescent, 48-hour release from quiescence without thymidine, and 48-hour release from quiescence with thymidine. Each point indicates the average centromere intensity level for all centromeres in a single cell, adjusted for background. Intensity values were normalized to cycling. Red line represents the median. Points were aggregated from 3 replicates. n = 181, 301, 137, and 97 for cycling, quiescent, no thymidine, and thymidine, respectively. All pairwise comparisons were significant with p = 0.321 between cycling and no thymidine and p < 0.0001 for all other comparisons using unpaired t test with Welch’s correction. M. Graph showing CENP-T centromere intensity in cycling cells treated with or without 2 mM thymidine for 48 hours. Each point indicates the average centromere intensity level for all centromeres in a single cell, adjusted for background. Intensity values were normalized to cycling. Red line represents the median. Points were aggregated from 3 replicates. n = 894, 337 for control and thymidine respectively. **** represents p<0.0001. N. Graph showing CENP-O/P centromere intensity in cycling cells treated with or without 2 mM thymidine for 48 hours. Each point indicates the average centromere intensity level for all centromeres in a single cell, adjusted for background. Intensity values were normalized to cycling. Red line represents the median. Points were aggregated from 3 replicates. n = 422, 197 for control and thymidine respectively. **** represents p<0.0001.

## References

1 Marescal, O. & Cheeseman, I. M. Cellular Mechanisms and Regulation of Quiescence. Dev Cell 55, 259–271, doi:10.1016/j.devcel.2020.09.029 (2020).

2 Kang, J. et al. In vivo self-renewal and expansion of quiescent stem cells from a non-human primate. Nat Commun 16, 5370, doi:10.1038/s41467-025-58897-x (2025).

3 Peng, Y. et al. RhoA-mediated G(12)-G(13) signaling maintains muscle stem cell quiescence and prevents stem cell loss. Cell Discov 10, 76, doi:10.1038/s41421-024-00696-7 (2024).

4 Kim, J. & You, Y. J. Oocyte Quiescence: From Formation to Awakening. Endocrinology 163, doi:10.1210/endocr/bqac049 (2022).

5 de Morree, A. & Rando, T. A. Regulation of adult stem cell quiescence and its functions in the maintenance of tissue integrity. Nat Rev Mol Cell Biol 24, 334–354, doi:10.1038/s41580-022-00568-6 (2023).

6 Berasain, C. & Avila, M. A. Regulation of hepatocyte identity and quiescence. Cell Mol Life Sci 72, 3831–3851, doi:10.1007/s00018-015-1970-7 (2015).

7 Schwabe, R. F. & Brenner, D. A. Hepatic stellate cells: balancing homeostasis, hepatoprotection and fibrogenesis in health and disease. Nat Rev Gastroenterol Hepatol 22, 481–499, doi:10.1038/s41575-025-01068-6 (2025).

8 Lei, L. et al. The mouse Balbiani body regulates primary oocyte quiescence via RNA storage. Commun Biol 7, 1247, doi:10.1038/s42003-024-06900-4 (2024).

9 Zhao, T. et al. Epigenetic maintenance of adult neural stem cell quiescence in the mouse hippocampus via Setd1a. Nat Commun 15, 5674, doi:10.1038/s41467-024-50010-y (2024).

10 van Velthoven, C. T. J. & Rando, T. A. Stem Cell Quiescence: Dynamism, Restraint, and Cellular Idling. Cell Stem Cell 24, 213–225, doi:10.1016/j.stem.2019.01.001 (2019).

11 Urban, N., Blomfield, I. M. & Guillemot, F. Quiescence of Adult Mammalian Neural Stem Cells: A Highly Regulated Rest. Neuron 104, 834–848, doi:10.1016/j.neuron.2019.09.026 (2019).

12 Coller, H. A., Sang, L. & Roberts, J. M. A new description of cellular quiescence. PLoS Biol 4, e83, doi:10.1371/journal.pbio.0040083 (2006).

13 Mitra, M., Batista, S. L. & Coller, H. A. Transcription factor networks in cellular quiescence. Nat Cell Biol 27, 14–27, doi:10.1038/s41556-024-01582-w (2025).

14 Kang, S., Antoniewicz, M. R. & Hay, N. Metabolic and transcriptomic reprogramming during contact inhibition-induced quiescence is mediated by YAP-dependent and YAP-independent mechanisms. Nat Commun 15, 6777, doi:10.1038/s41467-024-51117-y (2024).

15 Du, Y., Gupta, P., Qin, S. & Sieber, M. The role of metabolism in cellular quiescence. J Cell Sci 136, doi:10.1242/jcs.260787 (2023).

16 Fukada, S. et al. Molecular signature of quiescent satellite cells in adult skeletal muscle. Stem Cells 25, 2448–2459, doi:10.1634/stemcells.2007-0019 (2007).

17 Cheung, T. H. & Rando, T. A. Molecular regulation of stem cell quiescence. Nat Rev Mol Cell Biol 14, 329–340, doi:10.1038/nrm3591 (2013).

18 Subramaniam, S. et al. Distinct transcriptional networks in quiescent myoblasts: a role for Wnt signaling in reversible vs. irreversible arrest. PLoS One 8, e65097, doi:10.1371/journal.pone.0065097 (2014).

19 Min, M. & Spencer, S. L. Spontaneously slow-cycling subpopulations of human cells originate from activation of stress-response pathways. PLoS Biol 17, e3000178, doi:10.1371/journal.pbio.3000178 (2019).

20 Johnson, E. L., Robinson, D. G. & Coller, H. A. Widespread changes in mRNA stability contribute to quiescence-specific gene expression patterns in a fibroblast model of quiescence. BMC Genomics 18, 123, doi:10.1186/s12864-017-3521-0 (2017).

21 Chapman, N. M., Boothby, M. R. & Chi, H. Metabolic coordination of T cell quiescence and activation. Nat Rev Immunol 20, 55–70, doi:10.1038/s41577-019-0203-y (2020).

22 Ho, T. T. et al. Autophagy maintains the metabolism and function of young and old stem cells. Nature 543, 205–210, doi:10.1038/nature21388 (2017).

23 Liang, R. et al. Restraining Lysosomal Activity Preserves Hematopoietic Stem Cell Quiescence and Potency. Cell Stem Cell 26, 359–376 e357, doi:10.1016/j.stem.2020.01.013 (2020).

24 Venugopal, N. et al. The primary cilium dampens proliferative signaling and represses a G2/M transcriptional network in quiescent myoblasts. BMC Mol Cell Biol 21, 25, doi:10.1186/s12860-020-00266-1 (2020).

25 Breslow, D. K. & Holland, A. J. Mechanism and Regulation of Centriole and Cilium Biogenesis. Annu Rev Biochem 88, 691–724, doi:10.1146/annurev-biochem-013118-111153 (2019).

26 Azizzanjani, M. O. et al. Synchronized temporal-spatial analysis via microscopy and phosphoproteomics (STAMP) of quiescence. Sci Adv 11, eadt9712, doi:10.1126/sciadv.adt9712 (2025).

27 Palla, A. R. et al. Primary cilia on muscle stem cells are critical to maintain regenerative capacity and are lost during aging. Nat Commun 13, 1439, doi:10.1038/s41467-022-29150-6 (2022).

28 Maeshima, K. et al. Cell-cycle-dependent dynamics of nuclear pores: pore-free islands and lamins. J Cell Sci 119, 4442–4451, doi:10.1242/jcs.03207 (2006).

29 Feldherr, C. M. & Akin, D. Signal-mediated nuclear transport in proliferating and growth-arrested BALB/c 3T3 cells. J Cell Biol 115, 933–939, doi:10.1083/jcb.115.4.933 (1991).

30 Feldherr, C. M. & Akin, D. Regulation of nuclear transport in proliferating and quiescent cells. Exp Cell Res 205, 179–186, doi:10.1006/excr.1993.1073 (1993).

31 Coller, H. A. The paradox of metabolism in quiescent stem cells. FEBS Lett 593, 2817–2839, doi:10.1002/1873-3468.13608 (2019).

32 Wang, Y., Barthez, M. & Chen, D. Mitochondrial regulation in stem cells. Trends Cell Biol 34, 685–694, doi:10.1016/j.tcb.2023.10.003 (2024).

33 Baker, N. et al. The mitochondrial protein OPA1 regulates the quiescent state of adult muscle stem cells. Cell Stem Cell 29, 1315–1332 e1319, doi:10.1016/j.stem.2022.07.010 (2022).

34 McKinley, K. L. & Cheeseman, I. M. The molecular basis for centromere identity and function. Nat Rev Mol Cell Biol 17, 16–29, doi:10.1038/nrm.2015.5 (2016).

35 Vafa, O. & Sullivan, K. F. Chromatin containing CENP-A and alpha-satellite DNA is a major component of the inner kinetochore plate. Curr Biol 7, 897–900, doi:10.1016/s0960-9822(06)00381-2 (1997).

36 Palmer, D. K., O’Day, K., Trong, H. L., Charbonneau, H. & Margolis, R. L. Purification of the centromere-specific protein CENP-A and demonstration that it is a distinctive histone. Proc Natl Acad Sci U S A 88, 3734–3738, doi:10.1073/pnas.88.9.3734 (1991).

37 Mendiburo, M. J., Padeken, J., Fulop, S., Schepers, A. & Heun, P. Drosophila CENH3 is sufficient for centromere formation. Science 334, 686–690, doi:10.1126/science.1206880 (2011).

38 Barnhart, M. C. et al. HJURP is a CENP-A chromatin assembly factor sufficient to form a functional de novo kinetochore. J Cell Biol 194, 229–243, doi:10.1083/jcb.201012017 (2011).

39 Stirpe, A. & Heun, P. The ins and outs of CENP-A: Chromatin dynamics of the centromere-specific histone. Semin Cell Dev Biol 135, 24–34, doi:10.1016/j.semcdb.2022.04.003 (2023).

40 Swartz, S. Z. et al. Quiescent Cells Actively Replenish CENP-A Nucleosomes to Maintain Centromere Identity and Proliferative Potential. Dev Cell 51, 35–48 e37, doi:10.1016/j.devcel.2019.07.016 (2019).

41 Cheeseman, I. M. & Desai, A. Molecular architecture of the kinetochore-microtubule interface. Nat Rev Mol Cell Biol 9, 33–46, doi:10.1038/nrm2310 (2008).

42 Nishino, T. et al. CENP-T-W-S-X forms a unique centromeric chromatin structure with a histone-like fold. Cell 148, 487–501, doi:10.1016/j.cell.2011.11.061 (2012).

43 McKinley, K. L. et al. The CENP-L-N Complex Forms a Critical Node in an Integrated Meshwork of Interactions at the Centromere-Kinetochore Interface. Mol Cell 60, 886–898, doi:10.1016/j.molcel.2015.10.027 (2015).

44 Hori, T. et al. CCAN makes multiple contacts with centromeric DNA to provide distinct pathways to the outer kinetochore. Cell 135, 1039–1052, doi:10.1016/j.cell.2008.10.019 (2008).

45 Sissoko, G. B., Tarasovetc, E. V., Marescal, O., Grishchuk, E. L. & Cheeseman, I. M. Higher-order protein assembly controls kinetochore formation. Nat Cell Biol 26, 45–56, doi:10.1038/s41556-023-01313-7 (2024).

46 Screpanti, E. et al. Direct binding of Cenp-C to the Mis12 complex joins the inner and outer kinetochore. Curr Biol 21, 391–398, doi:10.1016/j.cub.2010.12.039 (2011).

47 Przewloka, M. R. et al. CENP-C is a structural platform for kinetochore assembly. Curr Biol 21, 399–405, doi:10.1016/j.cub.2011.02.005 (2011).

48 Falk, S. J. et al. Chromosomes. CENP-C reshapes and stabilizes CENP-A nucleosomes at the centromere. Science 348, 699–703, doi:10.1126/science.1259308 (2015).

49 Stellfox, M. E., Bailey, A. O. & Foltz, D. R. Putting CENP-A in its place. Cell Mol Life Sci 70, 387–406, doi:10.1007/s00018-012-1048-8 (2013).

50 Moree, B., Meyer, C. B., Fuller, C. J. & Straight, A. F. CENP-C recruits M18BP1 to centromeres to promote CENP-A chromatin assembly. J Cell Biol 194, 855–871, doi:10.1083/jcb.201106079 (2011).

51 Westhorpe, F. G., Fuller, C. J. & Straight, A. F. A cell-free CENP-A assembly system defines the chromatin requirements for centromere maintenance. J Cell Biol 209, 789–801, doi:10.1083/jcb.201503132 (2015).

52 Los, G. V. et al. HaloTag: a novel protein labeling technology for cell imaging and protein analysis. ACS Chem Biol 3, 373–382, doi:10.1021/cb800025k (2008).

53 French, B. T., Westhorpe, F. G., Limouse, C. & Straight, A. F. Xenopus laevis M18BP1 Directly Binds Existing CENP-A Nucleosomes to Promote Centromeric Chromatin Assembly. Dev Cell 42, 190–199 e110, doi:10.1016/j.devcel.2017.06.021 (2017).

54 Hori, T. et al. Association of M18BP1/KNL2 with CENP-A Nucleosome Is Essential for Centromere Formation in Non-mammalian Vertebrates. Dev Cell 42, 181–189 e183, doi:10.1016/j.devcel.2017.06.019 (2017).

55 Bodor, D. L., Valente, L. P., Mata, J. F., Black, B. E. & Jansen, L. E. Assembly in G1 phase and long-term stability are unique intrinsic features of CENP-A nucleosomes. Mol Biol Cell 24, 923–932, doi:10.1091/mbc.E13-01-0034 (2013).

56 Jansen, L. E., Black, B. E., Foltz, D. R. & Cleveland, D. W. Propagation of centromeric chromatin requires exit from mitosis. J Cell Biol 176, 795–805, doi:10.1083/jcb.200701066 (2007).

57 Su, K. C. et al. A Regulatory Switch Alters Chromosome Motions at the Metaphase-to-Anaphase Transition. Cell Rep 17, 1728–1738, doi:10.1016/j.celrep.2016.10.046 (2016).

58 Pedeux, R. et al. Thymidine dinucleotides induce S phase cell cycle arrest in addition to increased melanogenesis in human melanocytes. J Invest Dermatol 111, 472–477, doi:10.1046/j.1523-1747.1998.00324.x (1998).

59 Wang, R. C. & Wang, Z. Synchronization of Cultured Cells to G1, S, G2, and M Phases by Double Thymidine Block. Methods Mol Biol 2579, 61–71, doi:10.1007/978-1-0716-2736-5_5 (2022).

60 Prendergast, L. et al. The CENP-T/-W complex is a binding partner of the histone chaperone FACT. Genes Dev 30, 1313–1326, doi:10.1101/gad.275073.115 (2016).

61 Carpenter, A. E. et al. CellProfiler: image analysis software for identifying and quantifying cell phenotypes. Genome Biol 7, R100, doi:10.1186/gb-2006-7-10-r100 (2006).

62 Gascoigne, K. E. et al. Induced ectopic kinetochore assembly bypasses the requirement for CENP-A nucleosomes. Cell 145, 410–422, doi:10.1016/j.cell.2011.03.031 (2011).

63 Dudka, D. et al. Adaptive evolution of CENP-T modulates centromere binding. Curr Biol 35, 1012–1022 e1015, doi:10.1016/j.cub.2025.01.017 (2025).

